# Cell line resources for the study of neurofibromin: functions, phenotypes, and drug discovery/development

**DOI:** 10.64898/2026.08.13.743550

**Authors:** Hui Liu, Jian Liu, Chao Li, Elena Luppi, Kimia Rayat-Sanati, Elias Awad, Erik Westin, David Bedwell, Matthew Hartman, Andre Leier, Corina Anastasaki, David H. Gutmann, Robert Kesterson, Deeann Wallis

## Abstract

Our labs have been studying neurofibromin function and phenotype for over a decade with the intent of generating targeted therapeutics for Neurofibromatosis type 1 (NF1). In the process, we have generated numerous human cell lines containing variants within the *NF1* gene. Herein, we present data characterizing these cell lines and make them publicly available for use by researchers both within and outside the NF1 community. We describe lines that contain both well-characterized patient-specific variants either at their endogenous locus or as exogenous cDNAs, as well as variants of uncertain significance (VUS), engineered as heterozygous, homozygous, and compound heterozygous variants. Methods to generate each line and subsequent validation steps are detailed including targeted sequencing, Western blot analysis for neurofibromin expression and ERK activation. The utility of each line is dependent on the variant of interest, the parental cell line, and the mechanism of action relevant to possible therapeutic targeting.

## INTRODUCTION

Neurofibromin (NF1) is a tumor suppressor best known for its role as a Ras-GTPase Activating Protein (GAP) inhibitor of RAS activity. Approximately 1 in 3000 individuals are born with germline pathogenic variants (PVs) within the *NF1* gene and have Neurofibromatosis type 1 (NF1), a neurogenic disorder primarily defined by its hallmark feature, the neurofibroma. However, loss of *NF1* gene function impacts many cell types to create a pleomorphic disorder^1^. Somatic loss of *NF1* is a known oncogenic driver and can be found in many cancers including malignant peripheral nerve sheath tumors (MPNST), glioma, breast, lung, leukemia, and melanoma^2^. The *NF1* gene is quite large, spanning 300 kb and has 57 constitutive exons and several alternatively spliced exons. In addition to its function inhibiting the RAS pathway, recent studies indicate it can directly interact with ERα as a transcriptional co-repressor. Neurofibromin can also directly interact with PD-L1 and prevent PD-L1 cell surface expression and secretion to modulate tumor immune responses^3,4^.

Cell line models are useful for scientific research as they provide a sustainable source of biological material that can be manipulated in a controlled environment at a relatively low cost. They are essential for studying the molecular mechanisms of disease and/or complex biological processes, and they are critical for developing and testing new drugs. Hence, developing both patient-derived or genetically engineered models has great utility.

Many cell lines for the study of NF1 already exist. There are numerous patient-derived cell lines including Schwann cells from tumors (MPNST, plexiform neurofibromas (pNF), and cutaneous neurofibromas (cNF)), brain tumors, fibroblasts, and iPSCs^5–8^. These have been critical to show how loss of *NF1* impacts neurogenesis and Schwann cell function. iPSCs are especially valuable as they can be differentiated into many cell types and/or used to create 3D organoids^9–14^. Many NF1 cell lines are available through NTAP and ATCC or by donation from other labs.

Despite these resources, new cell lines are essential to study allele-specific effects of a large number of patient variants or specific gene-targeted therapeutics owing to the type of genetic mutation. Thus, our lab has generated a number of cell lines that could be useful for the study of NF1. We have developed many new human cell lines that harbor either exogenous cDNAs or CRISPR edits of endogenous alleles; these include HEK293 cells, human Schwann cells, and iPSCs.

## RESULTS

### HEK293 cells containing CRISPR edits

HEK293 cells are well-characterized and have been utilized for decades as a human cell line that is easily cultured and transfected. Despite the “embryonic kidney” designation, these cells express numerous neuronal proteins and may originate from an immature neuronal lineage within the embryonic kidney culture^15^. They possess a complex karyotype, including 3-4 copies of Chromosome 17 (where *NF1* resides), largely due to the incorporation of human adenovirus DNA during transformation, contributing to its genetic instability and heterogeneity. Importantly, HEK293 cells express all Ras Isoforms and are reasonable avatars for studying neurofibromin function. We have used CRISPR engineering to target the *NF1* gene and generate patient specific variants. We have previously published an *NF1* null (-/-) cell line in which *NF1* contains frameshifts (c.86_111del, c.93_104del, and c.102_103ins)^16^ as well as additional cell lines with a cryptic splice variant within exon 17; c.1885G>A; p. G629R^17^. Due to our interest in modeling exon skipping approaches, we generated cell lines with variants in exons 47 (c.6948insT) (Fig 1A) and exon 52 (c.7648A>T; c.7643_7647delTGAGG; c.7642insA with c.7648A>T) (Fig 1B). After targeted sequencing around c.6948insT, only the variant of interest was identified and no wildtype (WT) alleles were detected. This is supported by Westerns indicating both complete loss of neurofibromin protein and increase in GTP-Ras signaling (p=0.009) and activated ERK levels (p=0.005). The recurrent, patient-specific c.7648A>T variant, results in a premature stop and splice acceptor r.7647_7675del29; pArg2550Stop. Although annotated as a nonsense variant, it leads to the creation of a new cryptic splice site that results in a 29 bp deletion and frameshift. After targeted sequencing, a clone with 3 different mutant *NF1* alleles was identified. Alleles include: 1) c.7648A>T; c.7638 C>T (to ablate PAM sequence); c.7656 A>T (creates AluI restriction site for screening); 2) c.del7643_7647; and 3) c.7648A>T; c.7642insA. All alleles lead to frameshifts and together result in complete loss of neurofibromin protein and increases in GTP-Ras (p=0.046) and elevated pERK (p=0.001).

**Figure 1:**
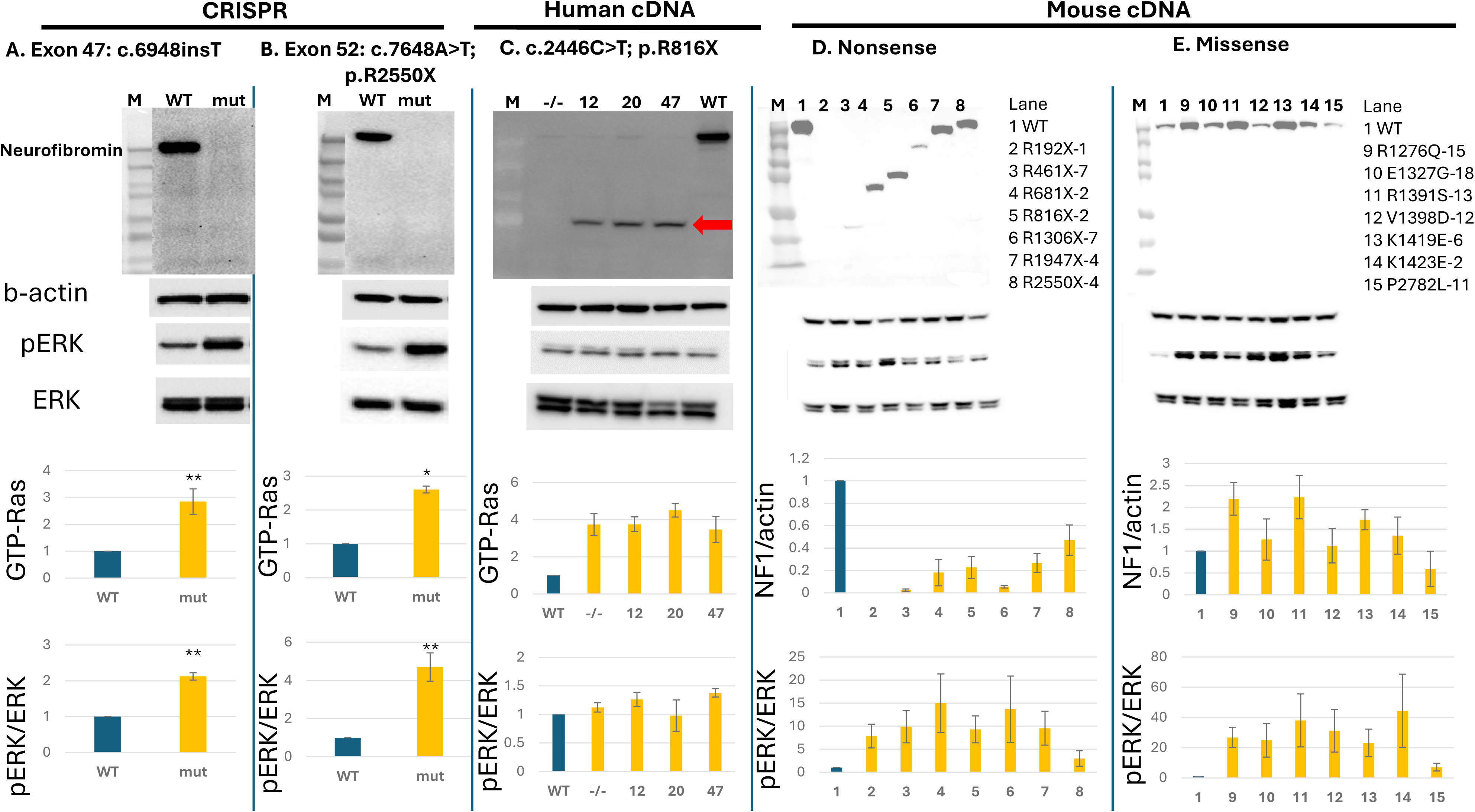
HEK2G3 cells containing endogenous CRISPR edits or expressing exogenous *NF1* cDNAs with variants. A-B. HEK293 cells containing endogenous CRISPR edits as indicated at top of each panel. From top to bottom: Representative Westerns for proteins as indicated along the left (neurofibromin, actin, pERK, ERK) showing complete loss of neurofibromin in mutant cell line. Histograms indicating GTP-Ras levels or pERK/ERK ratios (as a measure of ERK activation) of various cell lines C. HEK293 cells with exogenous human *NF1* cDNA containing R816X variant with bicistronic GFP. From top to bottom: Representative Westerns for proteins as indicated along the left (neurofibromin, actin, pERK, ERK) showing truncated neurofibromin (R816X) isoform as indicated by red arrow. Histograms indicating GTP-Ras levels or pERK/ERK ratios of various cell lines. The “-/-“ cell line is HEK293 cells containing inactivating indels in exon 2^16^. D. HEK293 cells with exogenous tags or PTCs. Representative Westerns showing truncated transcript sizes for neurofibromin along with actin, pERK, and total ERK. Histogram quantitating neurofibromin/actin levels in relation to the WT cDNA that has been tagged at the 3’ end. Histogram quantitating ERK activation in relation to the WT cDNA that has been tagged at the 3’ end E. HEK293 cell lines with expression of *NF1* cDNAs containing missense variants. Representative Western showing full length transcript sizes for all neurofibromin missense isoforms along with actin, pERK, and total ERK levels. Histogram quantitating neurofibromin/actin levels in relation to the WT cDNA that has been tagged at the 3’ end. Histogram quantitating ERK activation in relation to the WT cDNA that has been tagged at the 3’ end. *p<0.05 and **p<0.01 as determined by Student’s t-test.

#### HEK293 cells expressing exogenous *NF1* WT and variant cDNAs

We have published the development and utilization of expression plasmids containing full-length mouse *Nf1* cDNAs transiently transfected into HEK293 in which endogenous *NF1* has been deleted through CRISPR engineering (*NF1*^-/-^ cells) to help understand neurofibromin protein abundance (stability) and RAS inhibition^16,18^. As an extension of these studies, we have also generated stably transfected *NF1^-/-^* cell lines in which various human and mouse *NF1* cDNAs have randomly integrated into the HEK293 genome. We have used tagged cDNAs to affinity purify neurofibromin and its binding partners^19,20^. The tag we employed is inserted just 5’ of the stop codon in the cDNA and contains a TEV cleavage site and both a Strep II Tag and a 6XHis-Tag^19^.

As we are interested in nonsense suppression therapeutics (NSTs), we developed several HEK293 cell lines containing exogenous human *NF1* cDNA with variant R816X and bicistronic 2A-GFP cassette. We isolated multiple clones (12, 20, and 47) containing the cDNA. Western blots for neurofibromin (NF1) and actin reveal the truncated R816X protein (Fig 1C red arrow). While GTP-Ras levels remain elevated, activated ERK levels are not (Fig 1C).

We also generated a number of lines encoding premature termination codons (PTCs) in the murine *Nf1* cDNA, with PTCs including R192X, R461X, R681X, R816X, R1306X, R1947X, and R2550X variants (Fig 1D). These isoforms express truncated proteins in varying amounts as evidenced by lower molecular weight bands of variable intensity on Western blot. Differences in protein abundance likely reflect differences in genomic integration location, copy number, and intrinsic isoform stability. These truncated proteins are unable to completely normalize pERK/ERK ratios. Regardless, this can provide a good substrate to evaluate NSTs in terms of both increased full-length *NF1* transcript and decreased activated ERK in relation to untreated cells.

Additionally, we have generated a series of *NF1* missense variants within the GAP-related domain (GRD) to study the development of both Ras:neurofibromin stabilizers and neurofibromin mimetics. These GRD variants include R1276Q, E1327G, R1391S, V1398D, K1419E, and K1423E variants, as well as the VUS P2782L missense variant. All missense variants express full-length protein at varying levels that may depend on either its integration location within the genome, copy number, or the stability of the isoform. Regardless of protein abundance, none are able to fully normalize Ras-MAPK signaling, as reflected by elevated activated ERK.

### Schwann cells containing 3’ V5 tags on endogenous *NF1*

For patients with NF1-associated peripheral nerve sheath tumors, the most relevant cells are Schwann cells as these are the cell types that give rise to both cNFs and pNFs after undergoing loss of heterozygosity of *NF1*. Furthermore, pNFs may transform into MPNSTs. Hence, there is great interest in understanding the function of neurofibromin specifically in Schwann cells. Despite this demand, immortalized Schwann cell lines have been slow to become available, and patient derived immortalized lines have only been available for about a decade^5,6^. We utilized one of the immortalized WT cell lines ipn97.4 to generate an *NF1-*null cell line (termed ipn97.4^-/-^)^20^ and also utilized this parental WT line to “tag” the endogenous allele with a V5 epitope. Using CRISPR/Cas we created one cell line with V5 tags on both alleles (as confirmed by ddPCR and TOPO subcloning and sequencing of the 3’ end of *NF1*) called ipn97.4-V5-4 (Fig 2A). We also created a second cell line with a V5 tag on one allele and a 3’ indel that causes a frameshift before the terminal stop codon, termed ipn97.4-V5-82. Of note, despite the WT background of the tagged cell lines, the neurofibromin levels are in lower abundance (Fig 2A). This may be due to instability of the tagged allele that may get targeted to the proteosome for degradation. It is also worth noting that as anticipated, V5 is approximately twice as abundant in the cell line with both alleles tagged (4) as in the cell line with only a single allele tagged (82). Further, pERK/ERK ratios are not significantly different from WT levels.

**Figure 2:**
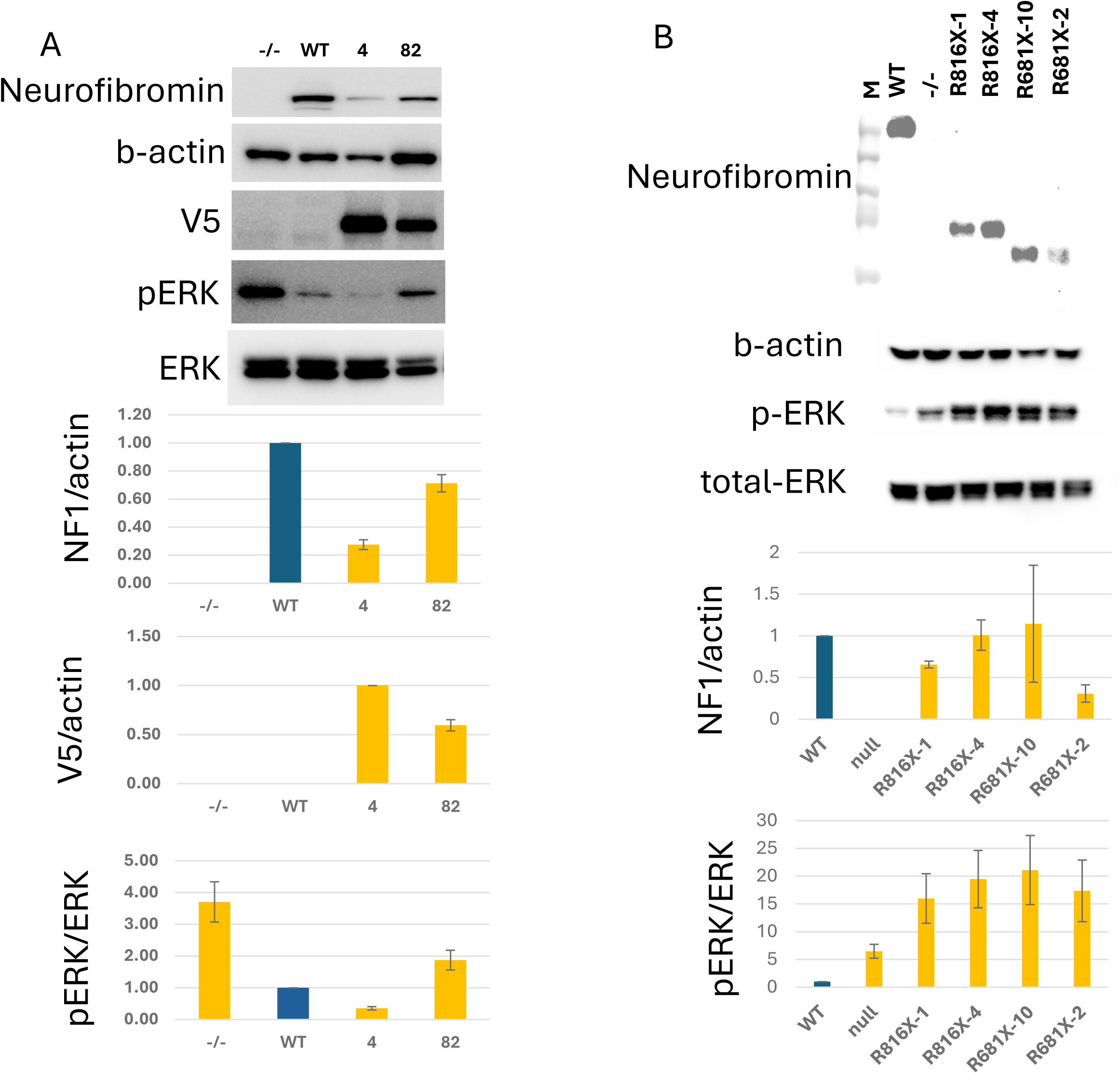
Schwann cells containing 3’ endogenous V5 tags or exogenous *NF1* cDNAs with PTCs R681X and R816X. A. Schwann cells containing 3’ endogenous V5 tags. Immortalized WT cell line ipn97.4 tagged on both alleles (ipn97.4-V5-4; or “4”) or a single allele (ipn97.4- V5-82, or “82”) with V5. From top to bottom: Representative Westerns of “-/-” (ipn97.4 NF1^-/-)^, “WT” (ipn97.4 NF1^+/+^) or tagged lines 4 and 82 of indicated proteins (neurofibromin, actin, V5, pERK and ERK. Histograms quantitating neurofibromin/actin, V5/actin, and pERK/ERK. B. *NF1* null Schwann cells containing exogenous *NF1* cDNAs with PTCs. From top to bottom: Representative Westerns of “WT” (ipn97.4; *NF1*^+/+^), “-/-” (ipn97.4; *NF1*^-/-^), and ipn97.4; *NF1*^-/-^ cells stably expressing *Nf1* cDNA with PTCs indicated. Histograms quantitating neurofibromin/actin and pERK/ERK levels in relation to WT.

### *NF1* null Schwann cells expressing exogenous *NF1* cDNAs with PTCs

We have also stably transfected our ipn97.4 *NF1^-/-^* Schwann cell line with various *Nf1* cDNAs. In the past, we used tagged cDNAs in Schwann cells to affinity purify neurofibromin and its binding partners^20^. We have also created null lines with stably transfected exogenous *NF1* cDNAs with PTCs including R681X and R816X (Fig 2B). These cell lines overexpress the variants of interest and the truncated neurofibromin protein is visible in varying levels of abundance on the Western blots (Fig 2B). Abundance may correlate with the integration site, copy number, or isoform stability. No full-length neurofibromin protein is detected by Western blotting. As anticipated, and as also seen in HEK293 cells (Fig 1D), these cDNAs are unable to normalize ERK activation.

#### iPSCs containing tags and CRISPR edits including PV and VUS

iPSCs are powerful models as they can be directed to differentiate into various cell types as well as into 3D organoid models. As such there has been great interest in developing these cell lines either through directed de-differentiation of patient derived cells or through CRISPR engineering^7^. We have CRISPR engineered multiple lines with tagged *NF1* alleles and several patient variants that include both PVs and VUS.

We generated various tagged alleles in which *NF1* has been endogenously tagged at the 3’ end. First, we created alleles that were tagged with a sensor tag to monitor protein abundance (HiBit) and/or an epitope tag (V5) (Figure 3A). These exist as compound heterozygous or homozygous tagged cell lines such that both *NF1* alleles are tagged. Cell lines were screened and validated with Westerns of: neurofibromin, actin (as loading control), HiBit, V5, pERK, and ERK (Figure 3B). It is notable that the 3’tags seem to reduce the abundance of neurofibromin protein expression to varying degrees. In certain cases, the lysines in the HiBit tag can be an alternate site of ubiquitination, and we suspect this may be a reason for the lower expression ^23^. Further, HiBiT has structured sequence and non-neutral biochemical properties, which may perturb protein folding or stability. In contrast, the V5 epitope is largely unstructured and inert.

**Figure 3:**
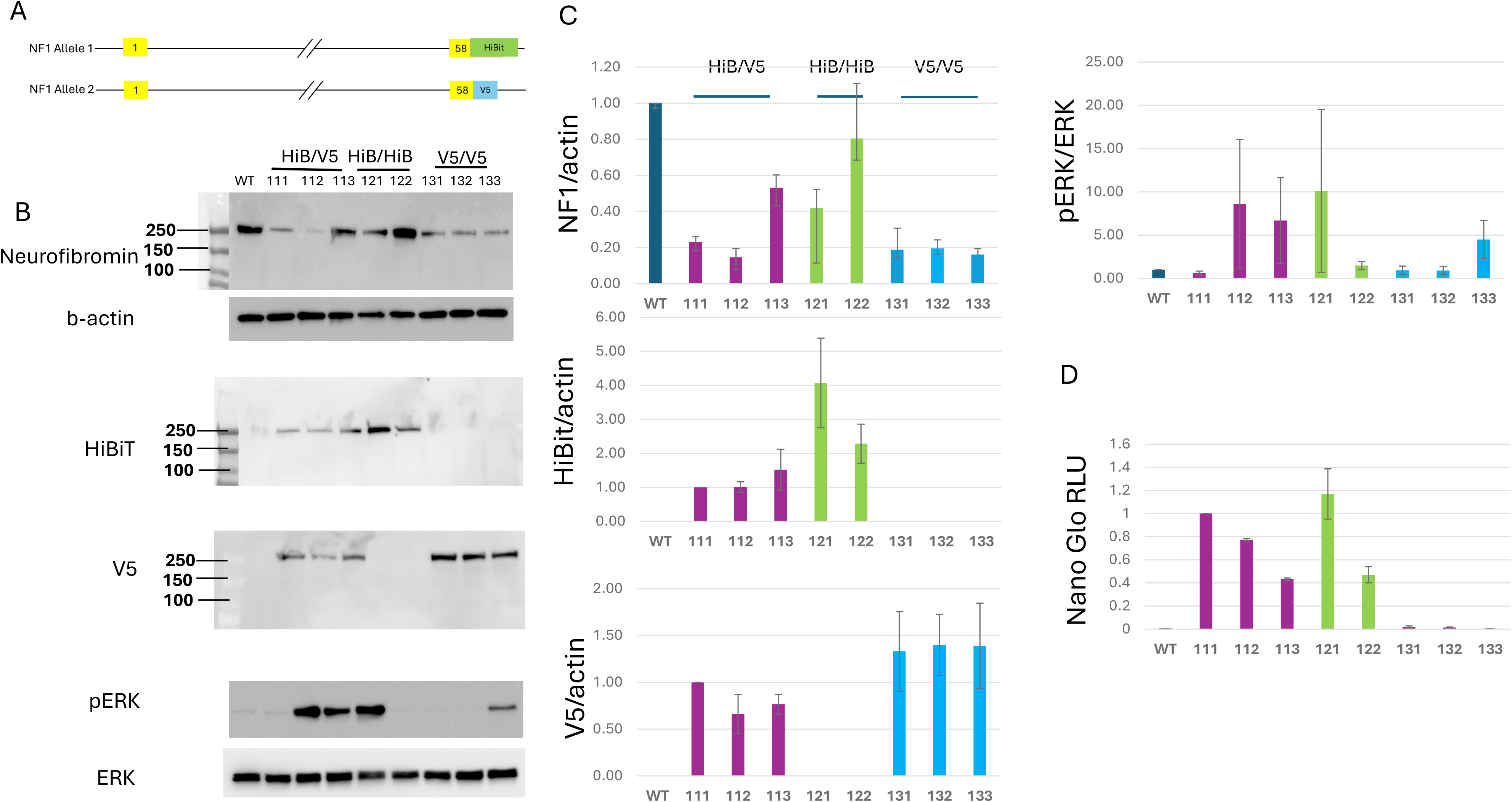
iPSC with WT *NF1* tagged at endogenous 3’ end with HiBit, V5, or both. A. Cartoon depicting 3’ *NF1* allele tags as compound heterozygotes. B. Representative Westerns of neurofiromin, actin, HiBit, V5, pERK and ERK for each clone identified. C. Histograms quantitating the Westerns are provided on the right (N=3 independent experiments). Dark blue bars indicate WT PGP1 cells without tags. Purple bars represent clones: NF-111, NF-112, and NF-113 which are compound heterozygous for the HiBit and V5 tags. Green bars represent clones NF-121 and NF-122 which have both alleles tagged with HiBit. Blue bars represent clones NF-131, NF-132, and NF-135 which have both alleles tagged with V5. D. Nano-Glo-HiBit lytic detection for each cell line.

Cells with two HiBit tags express relatively more HiBit protein via Western than cells with only one HiBit tag, though this does not translate to significantly higher activity in the Nano-Glo assay with HiBit lytic detection (Figure 3D); possibly due to variability within the assay. Also the cells with two V5 tags express relatively more V5 protein via Western than cells with only one V5 tag. We also see that activated ERK levels are extremely variable among these lines.

Next, we created *NF1* alleles that were tagged with HiBit-2A-Green Lantern (HiB-GL) and/or NanoLuc-2A-tdTomato (NLuc-tdT) (Figure 4A). These exist as compound heterozygous or homozygous tagged cell lines such that both *NF1* alleles are tagged. Cell lines were screened and validated with Westerns of: neurofibromin, GFP, HiBit, NanoLuc, TdTomato, actin, pERK, and ERK (Figure 4B). Note that NanoLuc protein expression is indicated with a red arrow. Westerns from two independent experiments were quantitated and presented as histograms for each antibody (Fig 4C). In addition, we used both Nano-Glo® HiBiT Lytic Detection System and Nano-Glo® Luciferase Assay System to detect HiBit and NLuc, respectively (Figure 4D). Finally, GFP and tdTomato levels were not readily detectable via fluorescent microscopy; thus, we used more sensitive FACs analysis to detect GFP and TdTomato signals in the appropriate cell lines (Figure 4E). We were able to detect modest levels of green in Green Lantern tagged cells in a dosage sensitive manner (Figure 4E (GFP mean fluorescent intensity (MFI)). We were also able to detect red in TdTomato in tagged cells in a dosage sensitive manner. It seems that cells with the HiB-GL tag express less neurofibromin protein than cells with only the NLuc-TdT. Despite the reduced neurofibromin expression, these cells have readily detectable GFP and HiBit expression by Western blotting. Cells expressing NLuc protein appear to do so in a dosage dependent fashion (e.g. cells with two copies express more than cells with a single copy). However, TdTomato protein does not appear to be dependent on neurofibromin expression dosage. Additionally, pERK is highly variable between the cell lines but is typically elevated in relation to the WT PGP1 cell line with no tags. Interestingly, while HiBit was easily detected by the Nano-Glo assay in the cells tagged with both HiBit-Green Lantern and Nluc-TdTomato; it was barely detected in cells containing two HiBit-Green Lantern alleles (Fig 4D). This is the opposite of what we might anticipate and may be due to the exceedingly low neurofibromin expression in these cells, possibly due to instability or degradation of the tagged and dimerized alleles. However, the NanoLuc-TdTomato allele is easily detectable with the Nano-Glo assay in cell lines with a single tagged allele and is about 2-fold higher in cell lines with both alleles tagged (Fig 4D).

**Figure 4:**
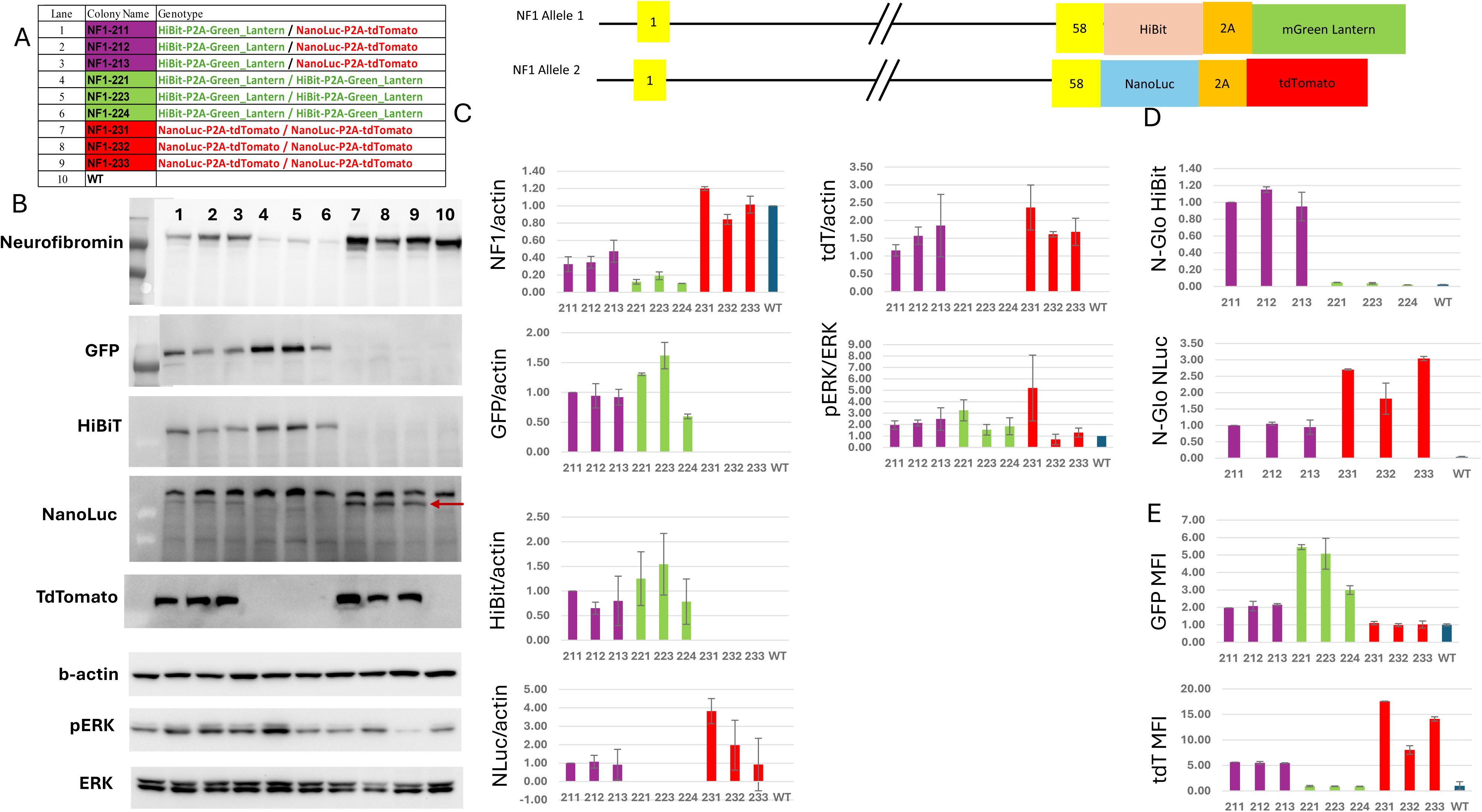
iPSC cells with WT *NF1* alleles with fluorescent tags at endogenous 3’ end. A. Table with Cell line (Colony) names (numbers) and genotypes that defines sample ID for subsequent validation and cartoon depicting *NF1* alleles with tag composition following exon 58. B. Representative Westerns for indicated cell lines and proteins (neurofibromin, GFP, HiBit, NanoLuc (red arrow), TdTomato, Actin, pERK and ERK. C. Histograms showing quantitation of N=2 Westerns for proteins indicated along y-axis. Dark blue bars indicate WT PGP1 iPSC line without tags. Purple bars represent clones: NF-211, NF-212, and NF-213 which are compound heterozygous for the HiBit-Green lantern and Nluc-TdTomato. Green bars represent clones NF-221 and NF-223, and NF-224 which have both alleles tagged with HiBit-Green Lantern. Red bars represent clones NF-231, NF-232, and NF-233 which have both alleles tagged with Nluc-TdTomato. D. Histograms for both HiBit and NanoLuc via Glo-assays from Promega (Nano-Glo HiBit and Nano-Glo Luc). E. FACs analysis for GFP MFI and TdTomato MFI.

Finally, we created iPSCs with patient specific variants introduced into the endogenous alleles using CRISPR Cas9. We have previously engineered an intragenic cryptic splice site (CSS) variant (c.1466A>G; p.Y489C) as a model for CSS suppression with antisense oligonucleotides^21^. In efforts to develop NSTs in an *NF1*-null background we utilized a patient derived iPSC heterozygous for variant c.2446C>T; p.R816X (generous gift from the Gutmann lab) and additional editing with CRISPR was used to generate a second variant allele. Targeted sequencing of the PCR product shows only the R816X variant. We evaluated the mutant allele frequency using ddPCR and found that while control iPSCs have 1% mutant allele frequency, both the heterozygous patient line and our clone of interest had 49% and 51% mutant allele frequency, supporting the idea that the targeted second *NF1* allele now contains an indel encompassing one or both PCR sequencing primers. We used Westerns to validate neurofibromin/actin and activated ERK (Fig 5A). Notably the heterozygous parental line expresses approximately 50% neurofibromin in relation to the WT PGP1 cell line and has approximately double the activated ERK (p=0.006). Once the second allele is removed, we see complete loss of neurofibromin protein, and much higher activated ERK (p=0.006).

**Figure 5:**
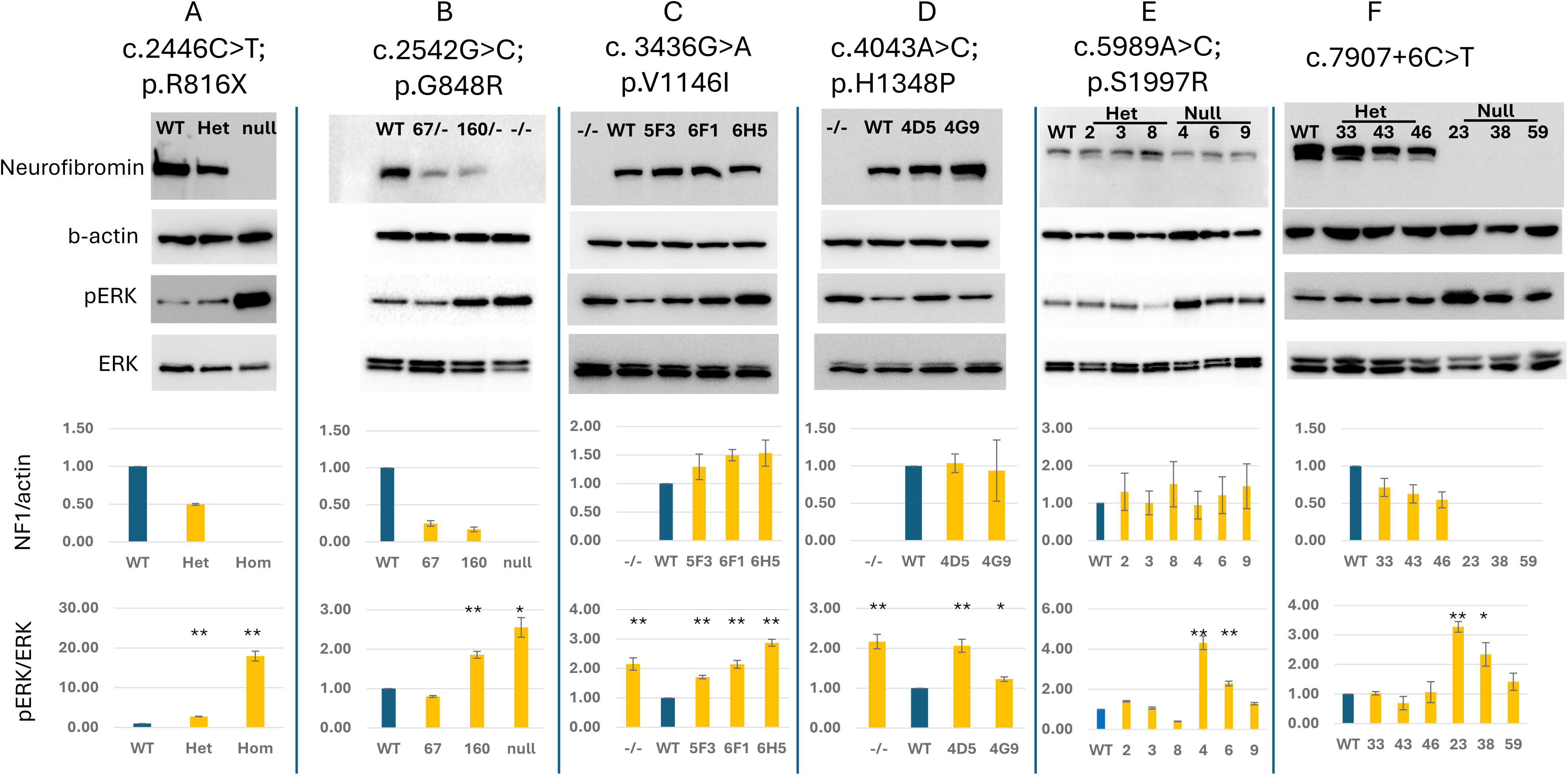
iPSC with patient specific variants introduced into the endogenous *NF1* alleles using CRISPR CasG. Variants are listed at the top of each panel. Cell lines were screened, sequenced, and validated by Western using indicated antibodies (neurofibromin, actin, pERK and ERK). Proteins were quantitated and represented below Westerns as histograms. A. c.2446C>T; p.R816X. WT represents PGP1 cell line, Het is a patient-derived iPSC containing R816X used for additional editing to generate “null” R816X with no neurofibromin expression and elevated pERK. B. c.2542G>C; p.G848R: WT represents PGP1 parental line, 67/- and 160/-represent two clones containing the G848R variant on one allele and an indel on the second *NF1* allele. -/- is a published iPSC line containing Y489C on both alleles. C. Clones containing c. 3436G>A p.V1146I in apparently homozygous state labeled as 5F3, 6F1, and 6H5; WT represents PGP1 parental line; and -/- is a published iPSC line containing Y489C on both alleles D. c.4043A>C; p.H1348P: WT represents PGP1 parental line; -/- is a published iPSC line containing Y489C on both alleles; 4D5 and 4G9 contain the c.4043A>C; p.H1348P VUS in apparently homozygous state. E. c.5989A>C; p.S1997R: WT represents the PGP1 parental cell line, Het indicates clones we identified containing the S1997R in heterozygous state and Hom indicates clones containing S1997R in homozygous state. F. 7907+6C>T: WT represents the PGP1 parental cell line, Het indicates clones we identified containing the c.7907+6T>G variant in heterozygous state and Hom indicates clones containing the variant in homozygous state. *p<.05, **p<.01 as determined by Student’s t-test.

To generate the c.2542G>C; p.G848R allele, we again used CRISPR and isolated and screened many clones. We were able to obtain compound heterozygous clones for this variant confirmed by targeted sequencing of PCR products. Both clones contain c.2542G>C; p.G848R on one allele and c.2559insG on the second allele. While the G848R isoform produces a full-length transcript based on Western blot analysis (Fig 5B), it is notable that these compound heterozygous lines do not produce 50% neurofibromin in comparison to WT cells. It has previously been reported that this allele may act in a dominant negative fashion upon dimerization^22^. ERK activation is also quite variable in these cell lines and only clone 160 has statistically elevated levels (p=0.003) (in addition to the -/- control (p=0.022)).

We generated several IPSC lines harboring VUS; c.3436G>A; p.V1146I, c.4043A>C; p.H1348P, c.5989A>C; S1997R, and c.7907+6T>G (a possible aberration of the canonical splice site).

First, we generated the c.3436G>A; p.V1146I variant which is located in the tubulin-binding domain of neurofibromin. Conflicting classifications have been reported in ClinVar (Variation ID: 141451; last accessed July 27, 2026), including 14 submissions classified as uncertain significance and 5 as likely benign, with the variant having been reported in both affected individuals and healthy populations. Out of 94 clones screened, we isolated and characterized three homozygous clones (5F3, 6F1, 6H5) via targeted Sanger Sequencing. No additional silent changes were introduced, as the variant itself disrupts the PAM sequence, thereby preventing Cas9 re-cutting after homologous repair. Neurofibromin protein levels are comparable to those observed in WT iPSCs (Fig 5C), indicating that the variant does not affect neurofibromin protein abundance. However, increased activated ERK, ranging from 1.71 to 2.87 fold compared to WT, indicates an impaired ability to regulate ERK activation (**p< 0.01 after Student’s t-test), suggesting that the variant may compromise NF1-mediated regulation of Ras signaling (Fig 5C).

Second, we generated c.4043A>C; p.H1348P, a variant located in the GRD domain of neurofibromin. This variant has been previously reported in ClinVar and classified as a VUS (Variation ID: 1737299; last accessed July 27, 2026). Out of 94 iPSC clones screened, 2 homozygous clones (4D5, 4G9) were generated, along with 3 additional silent variants introduced to promote homologous repair (c.4047C>T, c.4050C>T, c.4059C>T). In one of the clones (4G9), silent variant c.4059C>T is detected in heterozygosity, further supporting the homozygous genotype of the variant of interest. Here, we detected neurofibromin protein levels comparable to WT (Fig 5D), suggesting the variant does not significantly affect neurofibromin protein abundance. Conversely, the trend toward increased activated ERK (ranging from 1.23- to 2.06-fold relative to WT) indicates that the variant(s) may have an impaired ability to suppress RAS signaling (* p<0.05 or **p< 0.01 after Student’s t-test).

Third, iPSCs either heterozygous or homozygous for the c.5989A>C; S1997R VUS were generated and maintain neurofibromin protein levels (Fig 5E), but homozygous loss results in elevated ERK activation in some clones (**p< 0.01 after Student’s t-test). Heterozygous loss does not significantly increase ERK activation, though modest increases are evident. Published cDNA studies indicate that neurofibromin levels are relatively abundant at 91% of WT, but both GTP-Ras levels and ERK activation are increased suggesting S1997R remains stable, but is unable to fully repress Ras signaling activity in *NF1*-null cells^18^. Further, the Kesterson lab has generated a mouse model with this variant. As most *Nf1* PVs are embryonic lethal, we set heterozygous matings. After genotyping 53 pups from these crosses; there are 20 WT, 33 het, and 0 null. As normally we would expect a Mendelian ratio of 13.25: 26.5:13.25; Chi square = 18.3 with 2 degrees of freedom and p=.0001; indicating this is embryonic lethal and suggesting it is a pathogenic allele in mice.

Fourth, we used CRISPR to model a variant that might impact the canonical splice site; c.7907+6T>G. We were able to isolate multiple cell lines with c.7907+6T>G both as heterozygous and homozygous alleles out of 50 total clones screened. Heterozygous clones appear to express about 50% neurofibromin and do not have significantly elevated pERK, but homozygous clones do not express neurofibromin and have significantly elevated pERK (Figure 5F) (*p<0.05 and **p< 0.01 after Student’s t-test).

## Discussion

Herein we describe the development and preliminary characterization of a number of human cell lines that can be used to study NF1 and develop various types of therapeutics, paving the way for a better understanding of the mechanisms underlying phenotype modulation. One limitation of our characterization after CRISPR editing is that since we performed targeted sequencing, it is possible that one or more of the *NF1* alleles contains a deletion that encompasses the primer binding sites such that they are not amplified and sequenced. It is possible that these cell lines are indeed not homozygous null for the *NF1* variant of interest, but contain an additional, undetected deletion allele. For some of these cases, quantitative ddPCR analysis helped mitigate this limitation, while for others, indirect approaches, such as the detection of a heterozygous SNP near the desired edit and Western blot analysis of neurofibromin protein abundance can provide suggestive evidence. We were unable to further validate by ddPCR for the following cell lines: HEK293 Exon 47: c.6948insT, iPSC homozygous for c.5989A>C; p.S1997R (clones 4, 6 and 9); and iPSC homozygous for c.7907+6 T>G (clones 5, 9, and 17.)

We also see quite variable ERK activation between the cell lines, even when the variant(s) is the same. This has been our experience in almost all the cell lines we generate. For lines that undergo single cell cloning to derive a “pure” population, variation may be due to clone-to-clone variability. For lines containing integrated cDNA, variability may be the result of any combination of the following: integration site, copy number, isoform stability, or the drug selection process. Regardless, the variability in activated ERK within and between cell lines requires careful experimental design including careful cell culture handling and planning for control cell lines and treatment conditions.

While we quantitate and compare neurofibromin and ERK activation in cell lines with endogenous CRISPR variants, we do not try to directly compare cell lines with exogenous cDNAs or tags. For the exogenous cDNAs, our concern is that the expression levels may not reflect the biology of the variant as the genomic integration site or copy number may play a significant role. Similarly, for tagged proteins the inserted tags (with different lengths and 3D conformations) may interfere with stability of the resultant protein in ways that we are unable to predict at this time.

## Materials and Methods

### HEK293 cells containing CRISPR edits

HEK293 (WT or *NF1*^+/+^) cells were obtained from ATCC (CRL-1573) and cultured in DMEM + 10% FBS and 1X Pen/Strep using standard culture procedures. CRISPR guides were designed using CRISPOR (crispor.tefor.org), and a repair template was designed to generate a known patient mutation and also introduce silent changes to obliterate the protospacer adjacent motif (PAM) site and generate a restriction site for screening as necessary (Table 1). Guides were cloned and transfected into HEK293 cells. HEK293 cells were chosen because this cell line is well characterized, used historically in NF1 research, easily takes up exogenous DNA, and is easy to culture and scale. This cell line is derived from human embryonic kidney cells and carries a modal chromosome number of 64 in 30% of cells, and chromosome 17 (*NF1*) is present in 3–4 copies. Notably, this increased chromosome number does not affect any of the RAS or Ras-GAP genes. HEK293s have all three Ras isoforms. Cells were selected with puromycin, and single-cell cloning was used to isolate clonal lines for screening. DNA was isolated from each clone and screened for mutations of interest via restriction digest or direct sequencing of PCR products. PCR amplification primers are indicated in Table 1. Once clones of interest were identified, PCR products were cloned and individually sequenced to define alleles. Finally 1-3 clones containing the variant of interest were identified and further characterized via Western and or Ras-GLISA.

**Table 1:**
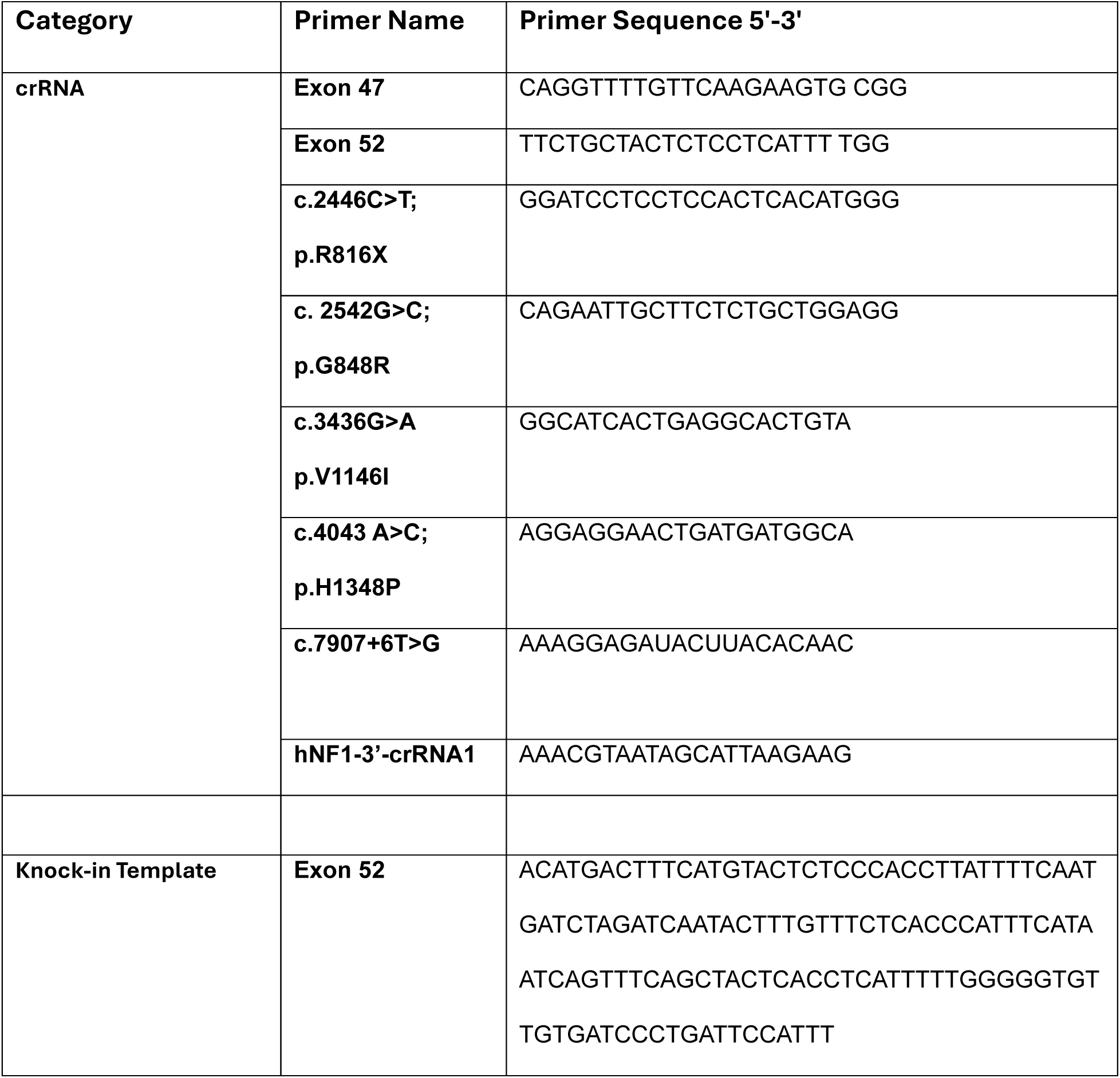
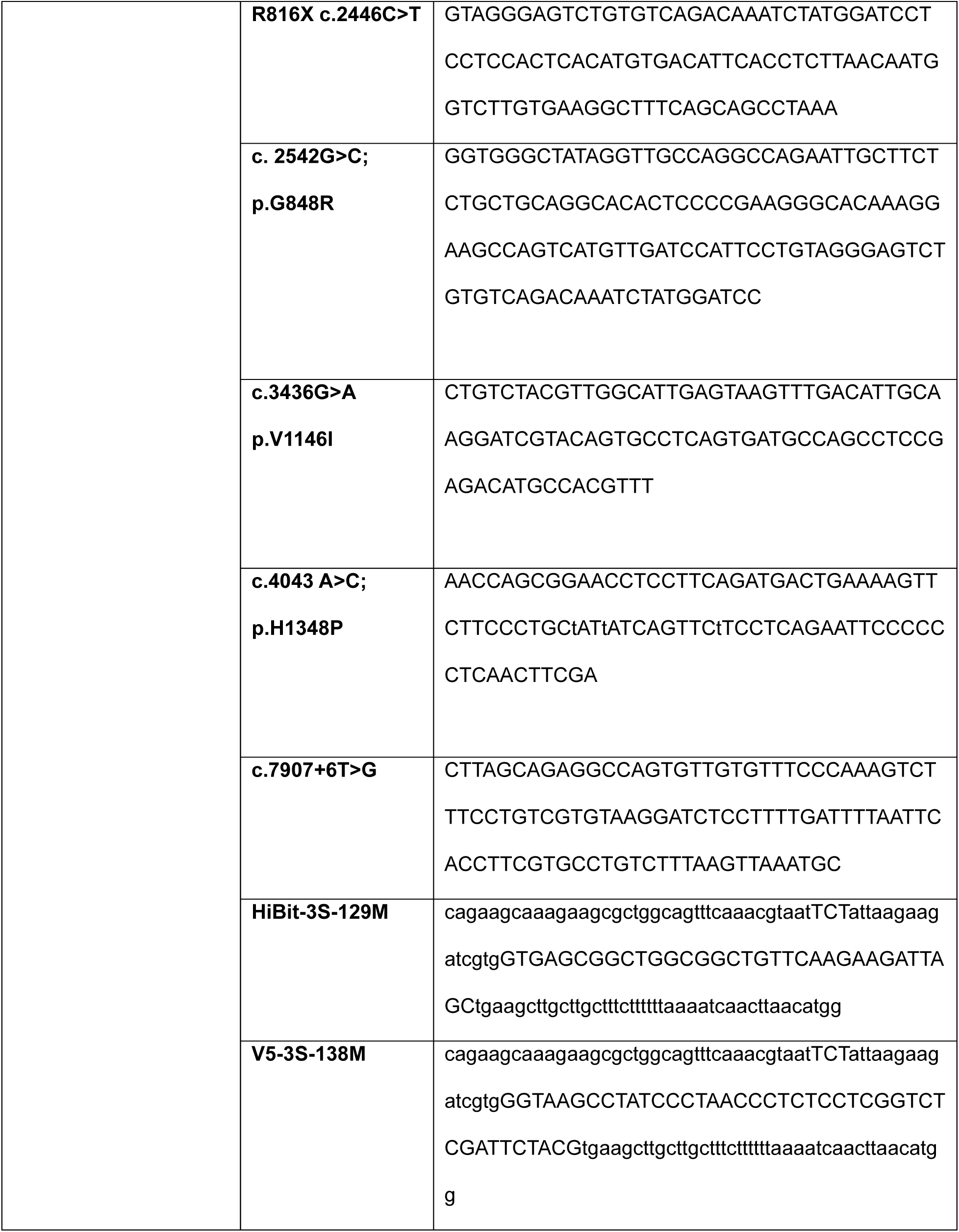
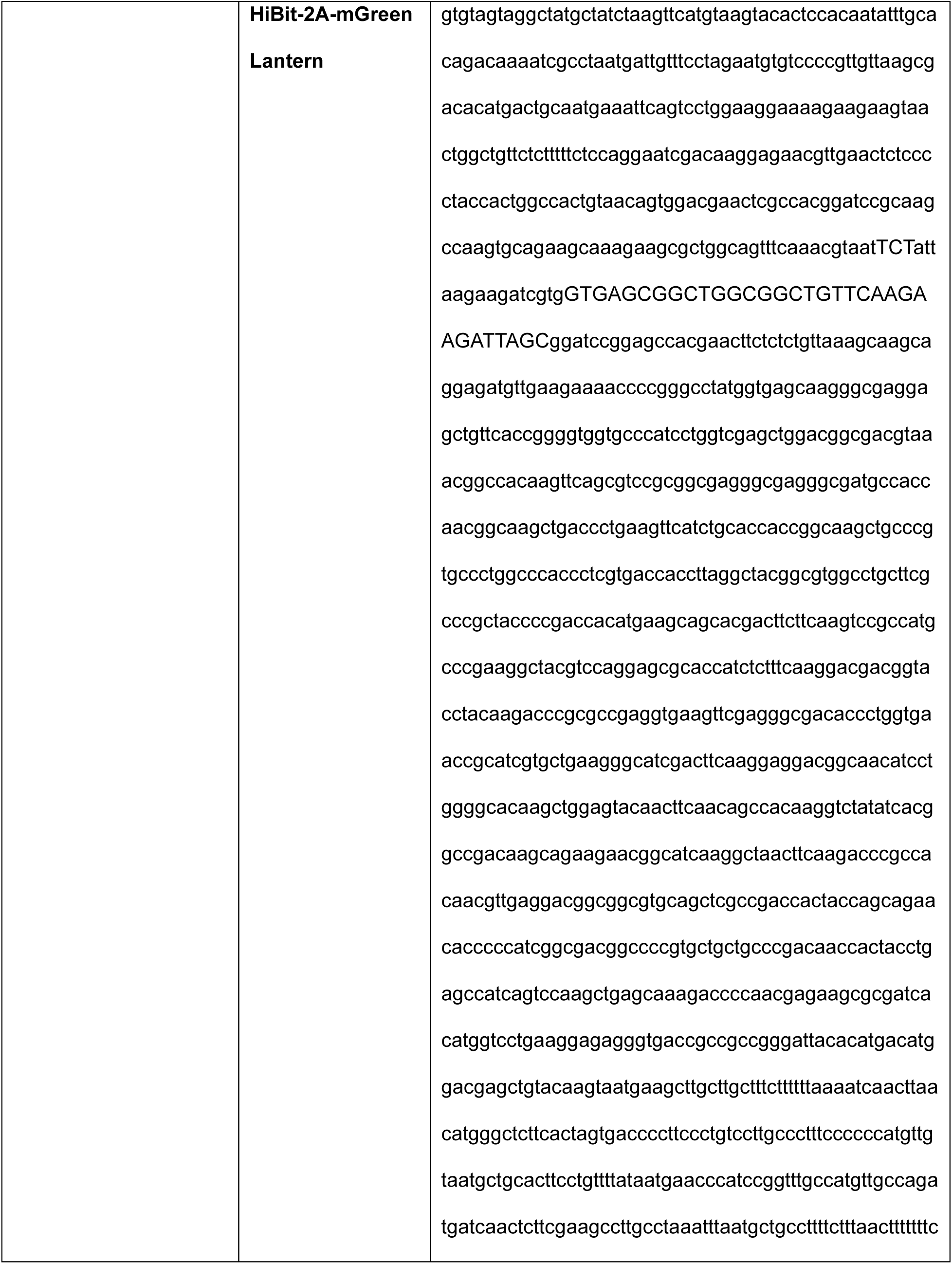
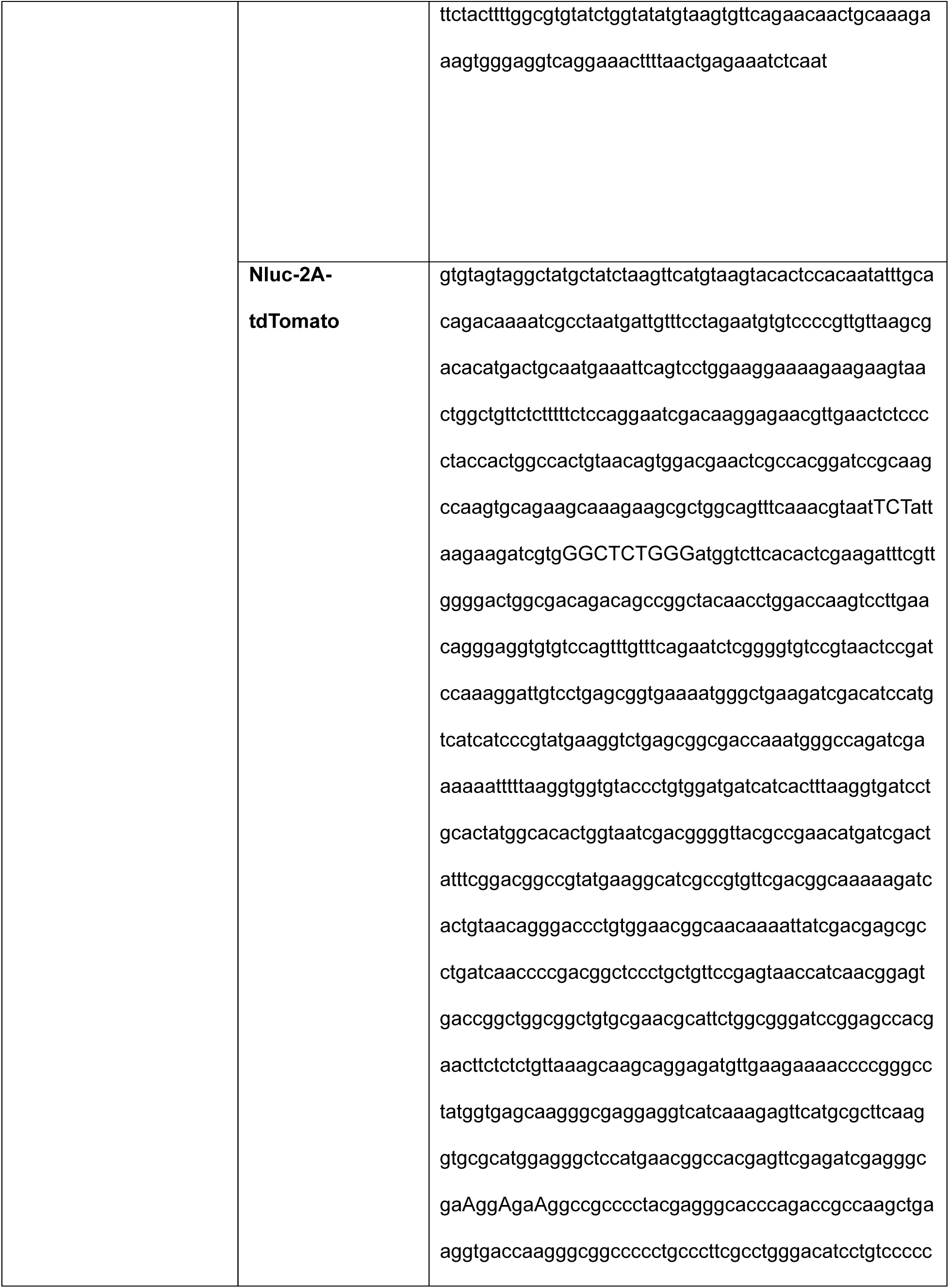
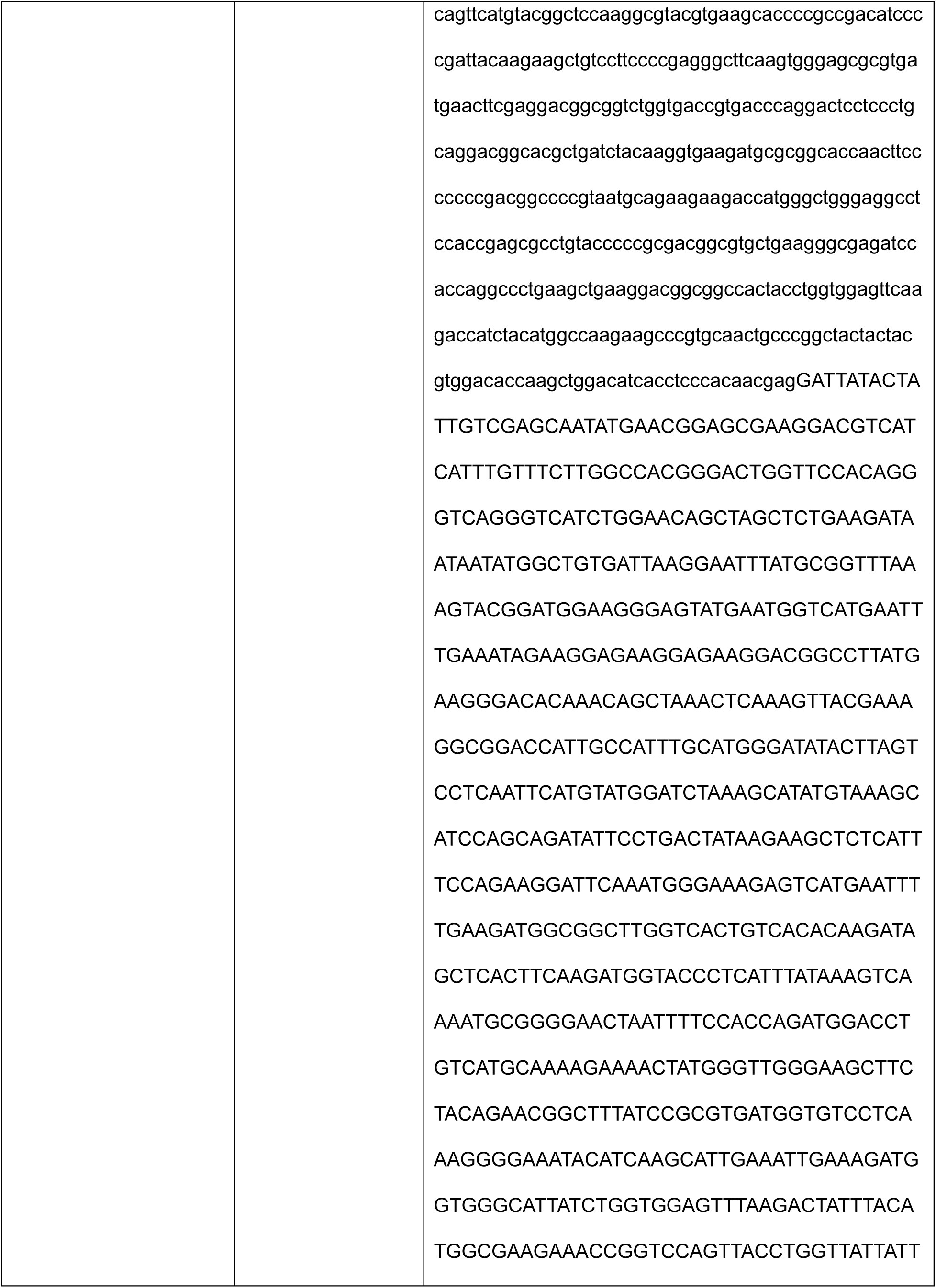
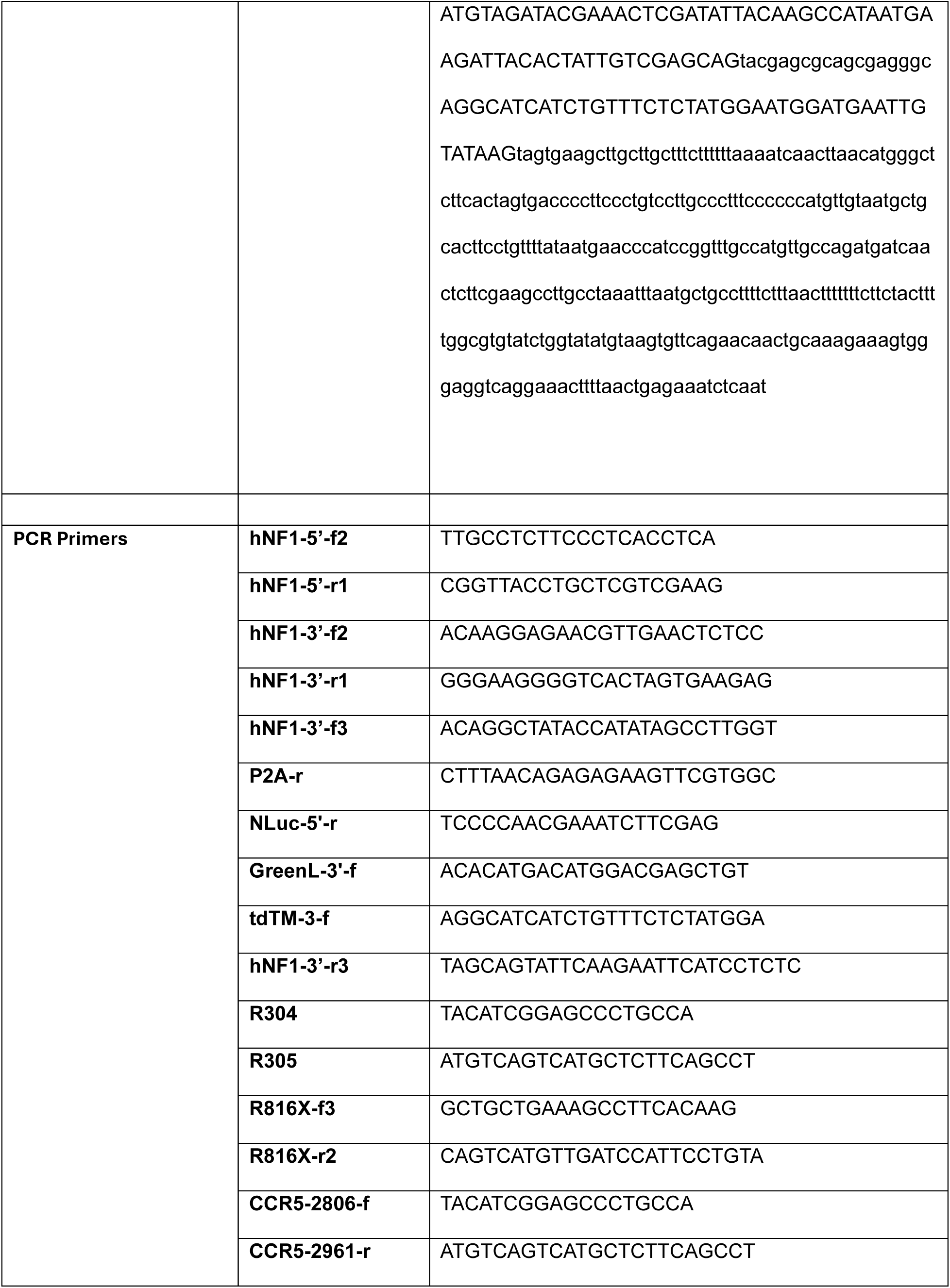
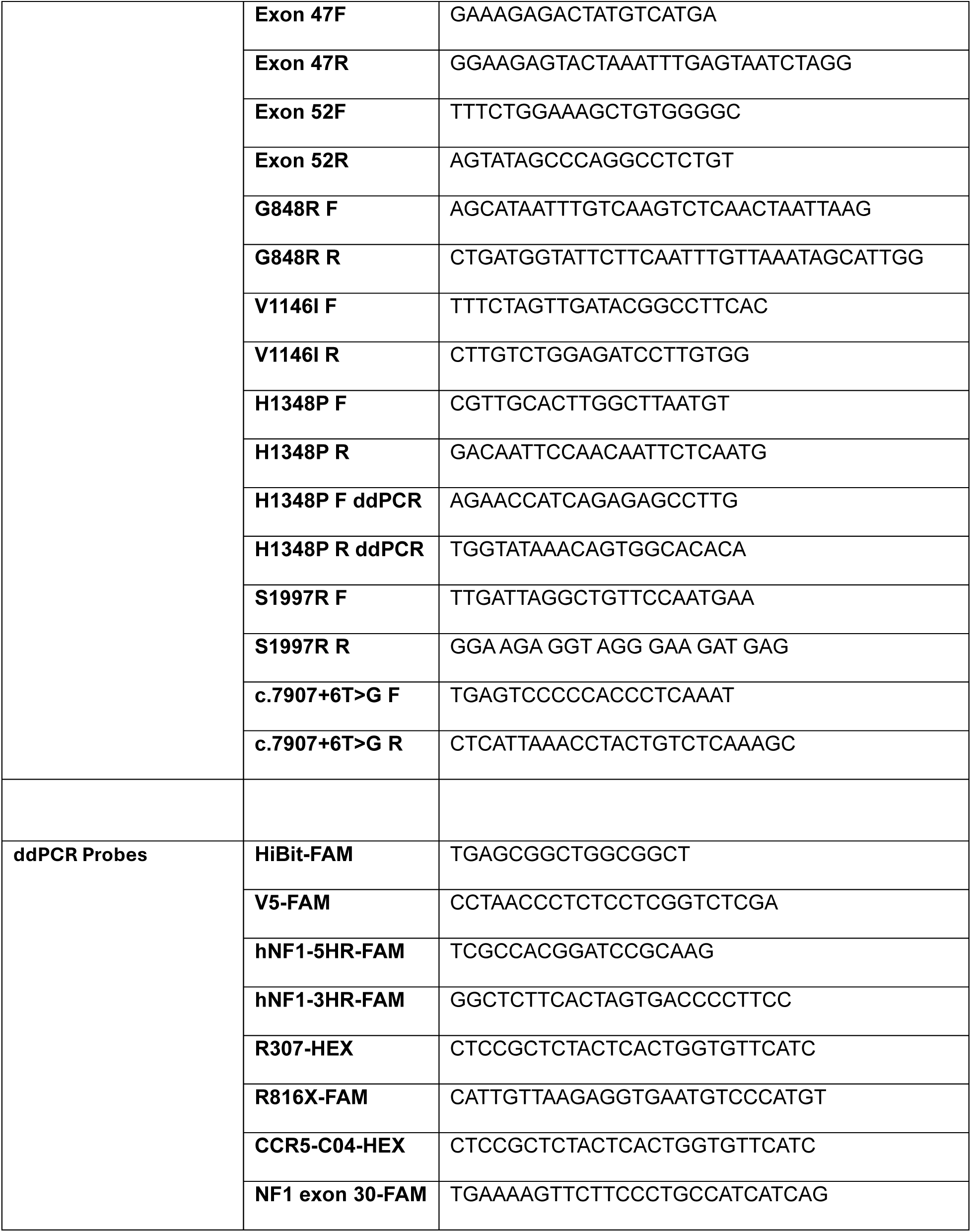
Guides, Repair Templates, Primers, and Probes.

### Western Blots

Protein lysates were harvested and quantified using a Bradford assay. Protein was utilized for Western blot and denatured at 70 °C for 10 min. Samples were loaded on Bio-Rad (Hercules, CA, USA) 4%–20% gradient gels (Cat. #: 4568094) and run at 100 V for 2 h. The gels were transferred onto a polyvinylidene difluoride (PVDF) membrane at 85 V for 2 h. Blots were probed overnight at 4 °C with selected primary antibodies for validation. Primary antibodies include neurofibromin antibody (Cell Signaling Technologies Cat. #: 14623S; Danvers, MA, USA), b-actin (Cell Signaling cat# 3700 1:1000), p-ERK (Cell Signaling cat# 9101 1:1000), total ERK (Cell Signaling cat# 9102 1:1000), V5 (Cell Signaling# 13202 1:1000), HiBit (Promega# N720A 1:1000), GFP (aveslabs # GFP-1020 1:1000), NanoLuc (Promega# N700A 1:1000) and TdT (Cell Signaling # 20163 1:1000). The next morning, the blots were washed and then probed with HRP tagged secondary from Santa Cruz. They were washed three more times before imaging using chemiluminescent substrate from BioRad (Cat. #: 170-5061; Hercules, California) as per the manufacturer’s protocols.

### RAS-G-LISA Assay

The RAS-G-LISA assay was obtained from Cytoskeleton Inc. and was performed according to the manufacturer’s instructions.

#### *NF1* null HEK293 cells and Schwann cells expressing exogenous *Nf1* cDNAs Stable transfection protocol

The murine *Nf1* cDNA development has been described^16,18^. We also developed a human *NF1* cDNA containing a 143 bp mini-intron between exons 35 and 36 under the control of CMV promoter with a 3’ 2A-GFP: pCEP4-CMV-NF1-2A-eGFP. This contains the EBNA-1 gene for episomal expression. Once an *NF1* wildtype clone was established we also introduced the R816X variant. The tagged cDNAs were transfected into either the *NF1* null HEK293 cells^16^ or *NF1* null Schwann cells^20^ as follows. Cells were seeded at 7.5 x 10^5^ cells per well in a 6 well plate. The following day HEK293 cells were transfected with 1ug of DNA and 3ul lipoD 293 reagent (Signagen Lab. Cat.SL100668), next day media was replaced with fresh media without transfection reagents or plasmids and cells recovered for 24h, then began with selection with 500 ug/ml G418 for 14-21 days. Schwann cells were transfected with 2.5 ug of DNA and 5 uL of Lipo 3000 reagent for 24h. The following day transfection with an additional 1 ug of DNA using LipoD while Lipo3000 mix is still in the wells. On the third day, media was replaced with fresh media without transfection reagents or plasmids and cells rested overnight. On the fourth day we began selection with 500 ug/ml G418 for 14-21 days until there were enough cells to pick colonies for single-cell selection. Cells remained under continuous selection while they were picked and harvested to test for neurofibromin protein expression via Western blot. Once confirmed, we continued to grow the cells in 10 cm dish with selection media until confluent and frozen back.

#### Establishment of HiBit and/or V5-tag -fused *NF1* knock-in induced pluripotent stem (iPS) cell lines and Schwann cells using CRISPR/Cas12a

PGP1 hiPSC cell line (RRID: CVCL_F182) was obtained from Synthego (Redwood City, CA, USA). All validation and QC were performed by the supplier. PGP1 (WT or *NF1* +/+) cells and their engineered derivatives were cultured in mTeSR Plus medium (Stemcell Technologies #100-0276) on Matrigel Basement Membrane coating Matrix (Corning #356234) in an incubator with humidified atmosphere at 5% CO2 and 37°C. When cells reached 70-80% confluency, they were routinely passaged using ReLeSR (Stemcell Technologies #100-0483) or EDTA. Briefly, cells were washed twice with PBS-EDTA medium (0.5 mM EDTA in PBS), then incubated with PBS-EDTA for 5 min at 37 °C. PBS-EDTA was removed, and cells were washed off swiftly with a small volume of corresponding medium.

To establish HiBit and/or V5-tag-fused *NF1* iPS and Schwann cells, our published protocol was adopted^24^. Briefly, 10^6^ cells were washed with DMEM/F12 medium (Thermo Fisher Scientific) and resuspended in 100 μl Nucleofector Solution (Lonza, P4 Primary Cell 4D-Nucleofector X Kit). Then, 160 pmol of Alt-R™ A.s. Cas12a (Cpf1) Ultra (IDT) was thoroughly mixed with 200 pmol hNF1-3’-crRNA1. The mixture was incubated at room temperature for 10 min. Then, 200 pmol ssODN (HiBit: HiBit-3S-129M, V5: V5-3S-138M) was mixed with Cas12a RNP. The final mixture was added into Nucleofector Solution containing the cells and gently mixed before electroporation (program CA167, Lonza 4D Nucleofector). Half the electroporated cells were cultured for 3 days and then genomic DNA was extracted. Gene targeting efficiency was examined by droplet-digital PCR (ddPCR) as follows: 11 μl 2× ddPCR Supermix for Probes (Bio-Rad), 1 μl hNF1-3’-f2 (20 μM), 1 μl hNF1-3’-r1 (20 μM), 1 μl HiBit-FAM (5 μM) or 1 μl V5-FAM (5 μM) , 1 μl R304 (20 μM), 1 μl R305 (20 μM), 1 μl R307-HEX (5 μM), 100 ng genomic DNA, add ddH2O to 22 μl of total volume). Then, 20 μl PCR mix was transferred to DG8 Cartridges (Bio-Rad) and droplets were generated with a QX200 Droplet Generator (Bio-Rad). Droplets were transferred into a ddPCR 96-Well Plate (Bio-rad), sealed with PX1 PCR Plate Sealer (Bio-Rad) and PCR was performed on a C1000 TouchThermal Cycler (Bio-Rad). The thermal cycling program conducted was step 1, 95 °C 10 min; step 2, 95 °C 30 s; step 3, 57 °C 3 min; repeat steps 2–3 39 times; step 4, 98 °C 10 min; and step 5, 8 °C hold. After PCR, droplets were analyzed using a QX200 Droplet Reader (Bio-Rad) with the ‘absolute quantification’ option. Once the percentage of HiBit or V5 gene targeting was identified, the other half of the electroporated iPS cells were used for isolating single-cell colonies as described^24^.

#### Establishment of HiBit-Green Lantern and/or NanoLuc-TdTomato-tag -fused *NF1* knock-in induced pluripotent stem (iPS) cell lines

Human iPSCs were maintained and passaged as described above. *NF1* knock-in iPSC lines carrying either a HiBit-Green Lantern or Nluc-TdTomato tag were generated using a CRISPR/Cas12a-based strategy adapted from the protocol described above. Briefly, 1×10^6^ iPSCs were resuspended in 100 μl Nucleofector Solution (Lonza, P4 Primary Cell 4D-Nucleofector X Kit) and electroporated with Cas12a ribonucleoprotein complexes assembled by incubating 160 pmol Alt-R A.s. Cas12a (Cpf1) Ultra (IDT) with 200 pmol hNF1-3’-crRNA1 for 10 min at room temperature. Electroporation was performed using program CA167 on a 4D-Nucleofector (Lonza). Recombinant AAV6 donor vectors carrying the *NF1* targeting cassette were produced in house by transfecting 293T cells seeded in five 150-mm dishes at 1×10^7^ cells per dish. After 24 h, each dish was transfected with 22 μg of the packaging/helper plasmid pDGM6 (a gift from D. Russell, University of Washington; Addgene plasmid 110660) and 6 μg of the transfer plasmid in 1 ml Opti-MEM I (Gibco, 31985088) using PEI (Polysciences, 23966-1). AAV6 was harvested 72 h after transfection and purified using an AAVpro Purification Kit (Takara, 6666) according to the manufacturer’s instructions. The donor cassette contained either the HiBit-Green Lantern or Nluc-TdTomato sequence flanked by *NF1* 5’ and 3’ homology arms. Immediately after electroporation, cells were plated onto Matrigel-coated plates in mTeSR1 supplemented with ROCK inhibitor, and the AAV6 donor was added shortly thereafter at an MOI of 1,000. Bulk knock-in efficiency was assessed 3 days after editing by genomic DNA extraction of half of the electroporated iPS cells followed by 5’ and 3’ junction ddPCR. For the *NF1* HiBit-Green Lantern donor, the 5’ assay used primers hNF1-3-f3 + P2A-r with probe hNF1-5HR-FAM, and the 3’ assay used primers GreenL-3’-f + hNF1-3-r3 with probe hNF1-3HR-FAM. For the *NF1* Nluc-TdTomato donor, the 5’ assay used primers hNF1-3-f3 + NLuc-5’-r with probe hNF1-5HR-FAM, and the 3’ assay used primers tdTM-3’-f + hNF1-3-r3 with probe hNF1-3HR-FAM. In all reactions, R304 + R305 and R307-HEX were used as the reference assay. ddPCR was performed using 2x ddPCR Supermix for Probes (Bio-Rad); droplets were generated with the QX200 Droplet Generator, amplified on a C1000 Touch Thermal Cycler using the following program: 95 °C for 10 min; 40 cycles of 95 °C for 30 s and 57 °C for 3 min; 98 °C for 10 min; and hold at 8 °C; and analyzed on a QX200 Droplet Reader using the absolute quantification setting. Once the percentage of HiBit-Green Lantern or Nluc-TdTomato gene targeting was identified, the other half of the electroporated iPS cells were used for isolating single-cell colonies as described^24^. Correctly targeted integration was defined by positive detection of both the 5’ and 3’ junctions.

#### Establishment of iPSC with CRISPR-induced *NF1* variants

To establish the c.2446C>T; p.R816X variant in conjunction with a null allele, we started with patient derived iPSC that were heterozygous for the allele. We then targeted using guides and repair templates detailed in Table 1. To generate G848R, we began with PGP1 cells and used detailed guides and repair templates.

We also applied CRISPR/Cas9 gene editing strategy to PGP1 cells to generate *NF1* VUS c.3436G>A; p.V1146I and c.4043A>C; p.H1348P models, using single guide molecules and repair templates designed through the Alt-R HDR Design Tool from IDT and listed in Table 1.

The RNP complex was assembled by incorporating 4μl of 100μM sgRNA and 6μl of 61μM HiFi Cas9 enzyme (IDT #1081060) per well at room temperature for 10 minutes. While the RNP complex was incubating, hiPSCs were dissociated into a single-cell solution using 1ml of Accutase (Stemcell #07920) and diluted to 1.000.000 cells/ml. The cells were spun down at 100rcf for 10 minutes and resuspended in 100 μl electroporation buffer (Lonza Nucleofector solution set P3 for primary cells #PBP3-00675) made of 82 μl nucleofection solution and 18 μL supplement solution.

Once incubation was completed, 10μl of RNP complex, 10 μl of ssHDR template and 100μl of cell suspension were mixed into a single Nucleocuvette. Then, the cell mixture was electroporated by Lonza 4D Nucleofector X Unit according to Pulse code CA167 for Primary cell P3. After electroporation, the cell mixture was left 10 minutes at RT and transferred into a single Matrigel-coated 60mm dish plate containing mTeSR Plus medium, ROCK inhibitor Y-27632 (Stemcell # 72304) at a final concentration of 10 μM and HDR Enhancer V2 at a final concentration of 1 μM (IDT #10007910). The plate was incubated in a tissue culture incubator (37°C, 5% CO2) for 72 hours allowing cell transfection. After the initial 24 hours, the culture medium was removed and replaced with fresh medium without HDR Enhancer and ROCK inhibitor. After 72 hours or when cells reached a suitable confluency, they were dissociated into a single-cell solution using Accutase, filtered through a 37μm reversible strainer, serial diluted to make the final concentration as 100 viable cells/ml and seeded one cell per well in a 96-well plate with 100 μL of mTeSR Plus medium containing CloneR2 (Stemcell # 100-0691) at 1X final concentration. The medium was replaced every other day, and the colonies were replica plated for DNA extraction and gene-editing screening via ddPCR/ restriction enzyme digestion when they occupied 70-80% of the well approximately after 10-12 days. CloneR2 was added only for the first 4 days. PCR primers and ddPCR probes are indicated in Table 1. Based on ddPCR results and/or restriction enzyme digestion patterns, clones were prioritized for genotype confirmation by targeted Sanger sequencing. Finally, 2-3 homozygous clones per variant were identified and further characterized.

ddPCR screening was performed for c.4043A>C variant as previously described with the following modifications. ddPCR Supermix for probes was used with 20 ng of genomic DNA. Probes and primers are described in Table 1. The WT probe was added to the final reaction mixture of 22 μl at a final concentration of 375 nM. The annealing temperature was set at 59.5C for 1 minute. Droplets were analyzed using a QX600 Droplet Reader (Bio-Rad) with the ‘direct quantification’ option.

To screen the CRISPR-mediated edits of interest, enzymatic digestion using the appropriate restriction endonuclease was also performed on PCR-products: Fatl (NEB #R0650) for c.4043A>C and AlwI (NEB #R0513) for c.3436G>A. PCR products were obtained using the Taq2x Master Mix (NEB #M0270) in a reaction mixture prepared in a final volume of 10μl and containing 20ng of genomic DNA. The primers are listed in Table 1. The amplification reaction was carried out using the T100™ Thermal Cycler instrument (Bio-rad), setting the following parameters: step 1, 95 °C 30 sec; step 2, 95 °C 15 s; step 3, 56 °C 30 sec; step 4 72 °C 30 sec; repeat steps 2–4 30 times; step 5 72 °C 5 min; and step 6, 8 °C hold. Each enzymatic digestion reaction was prepared in a total volume of 10 μl, composed of 1 μl target PCR product, 1 μl rCutSmart/r2.1 buffer (NEB #B6004), 0.5 μl restriction enzyme, 7.5 μl H_2_O. Samples were incubated at 37 °C for 1 hour. For each sample, an undigested control was included using the same reaction mix but omitting the enzyme. Following incubation, digested and undigested samples were loaded onto a 6% SDS–PAGE gel and electrophoresed at 150 V for 45–50 minutes. After electrophoresis, gels were stained in a 1× GelRed (Biotium #41002)/water solution for 5 minutes and imaged using a Bio-Rad ChemiDoc™ imaging system.

For VUS c.5989A>C; p.S1997R and c.7907+6T>G we utilized Synthego to generate knock-in pools of mutant iPSCs in the PGP1 parental background line. Wild type (WT) and variant PGP1 cells were sent to our lab for further single cell selection, screening and detection. We single cell cloned the pools of targeted cells and isolated single clonal cell lines with variants of interest. The variants were verified by sub-cloning and sequencing.

## Author Contributions

Chao Li, Methodology, Validation, Formal analysis, Investigation, Writing - Original Draft,

Hui Liu, Methodology, Validation, Formal analysis, Investigation, Data Curation, Writing - Original Draft

Jian Liu, Methodology, Validation, Formal analysis, Investigation, Data Curation, Writing - Original Draft

Elena Luppi, Methodology, Validation, Investigation, Data Curation, Writing - Original Draft

Kimia Rayat-Sanati, Validation, Formal analysis, Investigation, Elias Awad, Validation, Investigation, Writing - Review C Editing, David Bedwell, Writing - Review & Editing, Funding acquisition Matthew Hartman, Writing - Review & Editing, Funding acquisition Andre Leier, Writing - Review & Editing, Funding acquisition

Erik Westin, Methodology, Validation, Investigation, Data Curation

David Gutmann, Resources, Writing - Review C Editing,

Robert Kesterson Conceptualization Methodology, Writing - Review & Editing, Supervision, Project administration, Funding acquisition

Deeann Wallis Conceptualization Methodology, Formal analysis, Data Curation, Writing - Original Draft, Writing - Review & Editing, Visualization, Supervision, Project administration, Funding acquisition

## References

1 Staedtke, V. et al. Gene-targeted therapy for neurofibromatosis and schwannomatosis: The path to clinical trials. Clinical Trials 0, 17407745231207970, doi:10.1177/17407745231207970 (2023).

2 Philpott, C., Tovell, H., Frayling, I. M., Cooper, D. N. & Upadhyaya, M. The NF1 somatic mutational landscape in sporadic human cancers. Hum Genomics 11, 13, doi:10.1186/s40246-017-0109-3 (2017).

3 Berry, D. et al. The NF1 tumor suppressor regulates PD-L1 and immune evasion in melanoma. Cell Rep 44, 115365, doi:10.1016/j.celrep.2025.115365 (2025).

4 Zheng, Z. Y. et al. Neurofibromin Is an Estrogen Receptor-alpha Transcriptional Co- repressor in Breast Cancer. Cancer Cell 37, 387–402 e387, doi:10.1016/j.ccell.2020.02.003 (2020).

5 Li, H., Chang, L. J., Neubauer, D. R., Muir, D. F. & Wallace, M. R. Immortalization of human normal and NF1 neurofibroma Schwann cells. Lab Invest **96**, 1105–1115, doi:10.1038/labinvest.2016.88 (2016).

6 Li, H. et al. Immortalization and characterization of Schwann cell lines derived from NF1-associated cutaneous neurofibromas. PLoS One 21, e0340183, doi:10.1371/journal.pone.0340183 (2026).

7 Anastasaki, C. et al. Human iPSC-Derived Neurons and Cerebral Organoids Establish Differential Effects of Germline NF1 Gene Mutations. Stem Cell Reports 14, 541–550, doi:10.1016/j.stemcr.2020.03.007 (2020).

8 Anastasaki, C. et al. Aberrant coupling of glutamate and tyrosine kinase receptors enables neuronal control of brain-tumor growth. Neuron 113, 3582–3600.e3587, doi:10.1016/j.neuron.2025.08.005 (2025).

9 Xu, E. et al. Nested pediatric low-grade glioma cerebral organoid avatars reveal glutamatergic neuron stromal growth dependency. Genes Dev 40, 638–649, doi:10.1101/gad.353336.125 (2026).

10 Chen, A., Wang, H., Li, X., Anastasaki, C. & Gutmann, D. H. IRX2 and NPTX1 differential regulation of β-catenin underlies MEK-mediated proliferation in human neuroglial cells. Genes Dev **39**, 697–705, doi:10.1101/gad.352508.124 (2025).

11 Kuhrt, L. D. et al. Neurofibromin 1 mutations impair the function of human induced pluripotent stem cell-derived microglia. Dis Model Mech 16, doi:10.1242/dmm.049861 (2023).

12 Anastasaki, C. et al. Human induced pluripotent stem cell engineering establishes a humanized mouse platform for pediatric low-grade glioma modeling. Acta Neuropathol Commun 10, 120, doi:10.1186/s40478-022-01428-2 (2022).

13 Wegscheid, M. L. et al. Patient-derived iPSC-cerebral organoid modeling of the 17q11.2 microdeletion syndrome establishes CRLF3 as a critical regulator of neurogenesis. Cell Rep 36, 109315, doi:10.1016/j.celrep.2021.109315 (2021).

14 Mo, J. et al. Humanized neurofibroma model from induced pluripotent stem cells delineates tumor pathogenesis and developmental origins. J Clin Invest 131, doi:10.1172/jci139807 (2021).

15 Shaw, G., Morse, S., Ararat, M. & Graham, F. L. Preferential transformation of human neuronal cells by human adenoviruses and the origin of HEK 293 cells. FASEB J 16, 869–871, doi:10.1096/fj.01-0995fje (2002).

16 Wallis, D. et al. Neurofibromin (NF1) genetic variant structure-function analyses using a full-length mouse cDNA. Hum Mutat **39**, 816–821, doi:10.1002/humu.23421 (2018).

17 Leier, A. et al. Targeted exon skipping of NF1 exon 17 as a therapeutic for neurofibromatosis type I. Mol Ther Nucleic Acids 28, 261–278, doi:10.1016/j.omtn.2022.03.011 (2022).

18 Long, A. et al. Analysis of patient-specific NF1 variants leads to functional insights for Ras signaling that can impact personalized medicine. Hum Mutat, doi:10.1002/humu.24290 (2021).

19 Carnes, R. M., Kesterson, R. A., Korf, B. R., Mobley, J. A. & Wallis, D. Affinity purification of NF1 protein-protein interactors identifies keratins and neurofibromin itself as binding partners. Genes 10, 650 (2019).

20 Fay, C. X. et al. Global proteomics and affinity mass spectrometry analysis of human Schwann cells indicates that variation in and loss of neurofibromin (NF1) alters protein expression and cellular and mitochondrial metabolism. Scientific Reports 15, 3883, doi:10.1038/s41598-024-84493-y (2025).

21 Awad, E. K. et al. Restoration of Normal NF1 Function with Antisense Morpholino Treatment of Recurrent Pathogenic Patient-Specific Variant c.1466A>G; p.Y489C. J Pers Med 11, doi:10.3390/jpm11121320 (2021).

22 Young, L. C. et al. Destabilizing NF1 variants act in a dominant negative manner through neurofibromin dimerization. Proc Natl Acad Sci U S A 120, e2208960120, doi:10.1073/pnas.2208960120 (2023).

23 Lin, H. et al. Lysineless HiBiT and NanoLuc Tagging Systems as Alternative Tools for Monitoring Targeted Protein Degradation. ACS Medicinal Chemistry Letters 15, 1367–1375, doi:10.1021/acsmedchemlett.4c00271 (2024).

24 Li, C. et al. Novel HDAd/EBV Reprogramming Vector and Highly Efficient Ad/CRISPR-Cas Sickle Cell Disease Gene Correction. Sci Rep 6, 30422, doi:10.1038/srep30422 (2016).

